# No support for the effect of tDCS on frontal alpha asymmetry and behavioral and brain activity indices of inhibitory control

**DOI:** 10.1101/2023.09.20.558664

**Authors:** Atakan M. Akil, Renáta Cserjési, Tamás Nagy, Zsolt Demetrovics, Dezső Németh, H.N. Alexander Logemann

## Abstract

Prior research links self-regulation failure to psychiatric disorders. While the left and right dorsolateral prefrontal cortices are associated with distinct components of self-regulation—approach and inhibitory systems, respectively—existing measures are considered indirect. Our preregistered study explored frontal alpha asymmetry (FAA) as a potential biomarker for self-regulation. We explored whether the assumed effects of transcranial direct current stimulation (tDCS) on behavioral and brain activity indices of inhibitory control are mediated through changes in FAA. We used a randomized controlled sham-feedback design with 65 healthy humans (46 females). Before and after 2 mA anodal tDCS on the right frontal site, we collected resting-state EEG data to assess FAA scores, and participants also completed a stop signal task with neutral and intrinsic reward (food) conditions. The tDCS had no impact on FAA or any behavioral or neural indices of inhibitory control. However, event-related potential analyses revealed a correlation between inhibitory brain activity in the reward condition and trait FAA. Higher right relative to left frontal brain activity was linked to lower early-onset inhibitory activity, possibly originating from the inferior Frontal Gyrus, but correlated with higher late-onset inhibitory control, presumably originating from the superior Frontal Gyrus. While tDCS yielded unexpected outcomes in FAA and inhibitory control metrics, we found an interesting dissociation regarding the lateralization of frontal brain activity and early and late onset inhibitory brain activity.

## 1. Introduction

Self-regulation is the self’s ability to control its own thoughts, behaviors, and emotions ^1^, and its failure has been linked with various mental health problems such as substance abuse (e.g., cocaine ^2^, nicotine ^3^, alcohol ^4^, obesity ^5^, major depressive disorder ^6^, and internet addiction ^7^. A potential neuromarker and treatment target for psychiatric disorders associated with impaired self-regulation can be frontal alpha asymmetry (FAA), which is an electrophysiological (EEG) measure of the difference in alpha power (8-12 Hz) between the left and right dorsolateral prefrontal cortex (DLPFC). More specifically, self-regulation is driven by two main motivation systems, both of which are influenced by environmental factors ^8,9^: Environmental stimuli that indicate potential rewards trigger approach motivation, whereas stimuli signaling threat activate inhibitory motivation. Previous research found that these two antagonistic components may be associated with the asymmetry of frontal cortical activity ^9–11^: Relative left frontal cortical activity dominance is linked with approach tendency while right frontal cortical activity dominance is associated with inhibitory control. Here, our aim is to overcome the limitation of correlational studies by utilizing non-invasive brain stimulation to investigate frontal brain activity asymmetry and inhibitory skills in intrinsic reward contexts.

Inhibitory control, the ability to suppress an intended response, stands as a fundamental facet of self-regulation and relevant clinical conditions ^12^. Individual differences in inhibitory control are frequently assessed using the Stop Signal Task (SST) ^13^. In this task, participants are presented with “go” stimuli (typically symbols in visual tasks) at the center of the screen, requiring a button-press response. Occasionally, these go stimuli are followed by a “stop” stimulus (a specific symbol), signaling participants to withhold their response. The stop-signal reaction time (SSRT), a widely accepted measure of inhibitory control ^12,14^, is derived from this task.

To explore brain mechanisms underlying inhibitory control, prior studies have utilized a combination of the SST and event-related potential (ERP) analyses ^15,16^. ERPs reflect the synchronous activation of large groups of neurons time-locked to an event ^17^. The Stop N2, occurring around 200 ms latency, exhibits significantly more negative amplitudes during successful stop trials than unsuccessful ones ^13^. Furthermore, it has been implicated that the neurobiological correlate of the Stop N2 is the right inferior frontal gyrus ^13^. The Stop P3 is influenced by stopping success, showing larger amplitudes for successful inhibitions compared to failed ones ^13^. Indeed, the Stop P3 has been thought to represent inhibitory control ^18^ and to originate from the superior frontal gyrus ^19^. Despite these presumed associations, inhibitory control’s behavioral and EEG indices have not been explored in the context of FAA.

Transcranial direct current stimulation (tDCS) is a non-invasive neuromodulation technique that delivers electrical currents to the brain through electrodes placed on the scalp to modulate specific brain activities ^20^. In the context of our interests, promisingly, various studies have provided some support indicating that tDCS over DLPFC reduces cravings, desire to binge eat, and overall food intake. ^21–23^. However, even though there is evidence regarding the effects of tDCS over DLPFC on inhibitory control, these measures are indirect, and to the best of our knowledge, it is still unknown whether these tDCS-associated effects can be attributable to changes in resting-state FAA.

In this preregistered study, first, we hypothesized that an active tDCS (anode over right DLPFC/cathode over left DLPFC) would result in an increase in right relative to left frontal brain activity. Since the amplitude within the alpha frequency band represents inactivity, lower FAA scores (right minus left alpha) reflect relatively less left than right cortical activity ^10^; therefore, we expected to observe decreased FAA scores at frontal sites (F4-F3). Additionally, we anticipated that the application of active tDCS would result in increased inhibitory control within the food reward context, as opposed to the neutral context. This increase would manifest as a decreased SSRT in the SST, along with an increase in the Stop N2 and Stop P3.

## 2. Methods and Materials

### 2.1. Participants

Our pilot study (N = 10) regarding the main effect of tDCS on the primary outcome, FAA, served as the foundation of the sample size rationale. More specifically, the expected smallest effect (f > 0.237; η2partial > 0.053) was estimated to be detectable with the a priori intended sample size of 30 for the active intervention group using G*Power ^24,25^, with a defined power of 80 percent, alpha at .05, and estimated FAA test-retest correlation of 0.6. Sixty-five healthy humans (46 females), ranging in age from 18 to 58 (M_age_ = 23.93; SD_age_ = 6.08), were ultimately recruited via social media and university courses for the study after several participants terminated the experiment for a variety of reasons, such as boredom and fatigue. Each participant was given a voucher or course credit for their participation. In order to participate in the study, individuals needed to meet certain eligibility criteria. Specifically, they had to be at least 18 years of age. Additionally, participants were screened for certain exclusion criteria, which included the presence of psychological or psychiatric disorders, frequent headaches or migraines, metallic implants, epilepsy, significant head trauma in the past, recent head trauma, pacemaker usage, chronic skin conditions, and current drug use. Participants were also required to self-assess their proficiency in English using the Common European Framework of Reference for Languages - Self-assessment Grid ^26^, and a minimum of B1 level English proficiency was necessary for inclusion in the analyses. Psytoolkit was used for self-report assessments ^27,28^. Before any procedures took place, all participants provided their written informed consent. The research was conducted following the ethical guidelines outlined in the Declaration of Helsinki and its later amendments. The study was approved by the local research ethics committee.

### 2.2. Stop signal task (SST)

The SST was developed using OpenSesame ^29^. It was adapted from the original version ^13^ and included a food reward condition based on previous studies ^30,31^. The task consisted of 10 blocks in total, with one practice and four main blocks for each condition (neutral/reward) separately. Participants were seated approximately 65 cm. from the screen. At the beginning of the task, participants were instructed with an understandable text, and it was also explained orally in case it was needed. First, a fixation dot was displayed in the middle of the screen to engage attention and eye fixation for 2000 ms. Slightly above the dot, in the reward condition, palatable food pictures, i.e., cookies, chips, chocolate, and nuts (115 px width (2.9°) x 200 px height (5.1°)), which require a key response were presented randomly in horizontal or vertical orientation for 150 ms (Figure 1). Therefore, the reward condition had two dimensions (horizontal/vertical). On the other hand, in the neutral condition, instead of the palatable food pictures, the letters “X” or “O” (200 px width (5.1°) x 200 px height (5.1°)) were presented for 150 ms, which also requires a key response depending on the letter presented. Either a left-button press or a right-button press with the left and right index fingers based on a stimulus was used for responses. In stop trials, go stimuli (either palatable food pictures in horizontal or vertical position or letters X or O depending on the condition) were infrequently followed by the letter “S” (200 px width (5.1°) x 200 px height (5.1°)) for 150 ms representing “stop” and participants were supposed to withhold their response in that case. The go-stop Stimulus Onset Asynchrony (SOA) was fixed at 350 ms, but when participants could not withhold their response, the timing was arranged after each stop trial via an algorithm; the go-stop interval was decreased 50 ms. In case of a successful inhibition, the interval increased 50 ms. The tracking algorithm yields an approximate 50% inhibit rate, increasing the reliability of the estimation of the SSRT ^32^. In total, a session of the task took approximately 45 minutes. The main outcome of the SST was the SSRT. By following Verbruggen et al. ^32^, omissions were replaced with maximum reaction times (RTs) (1500 ms).

**Figure 1.**
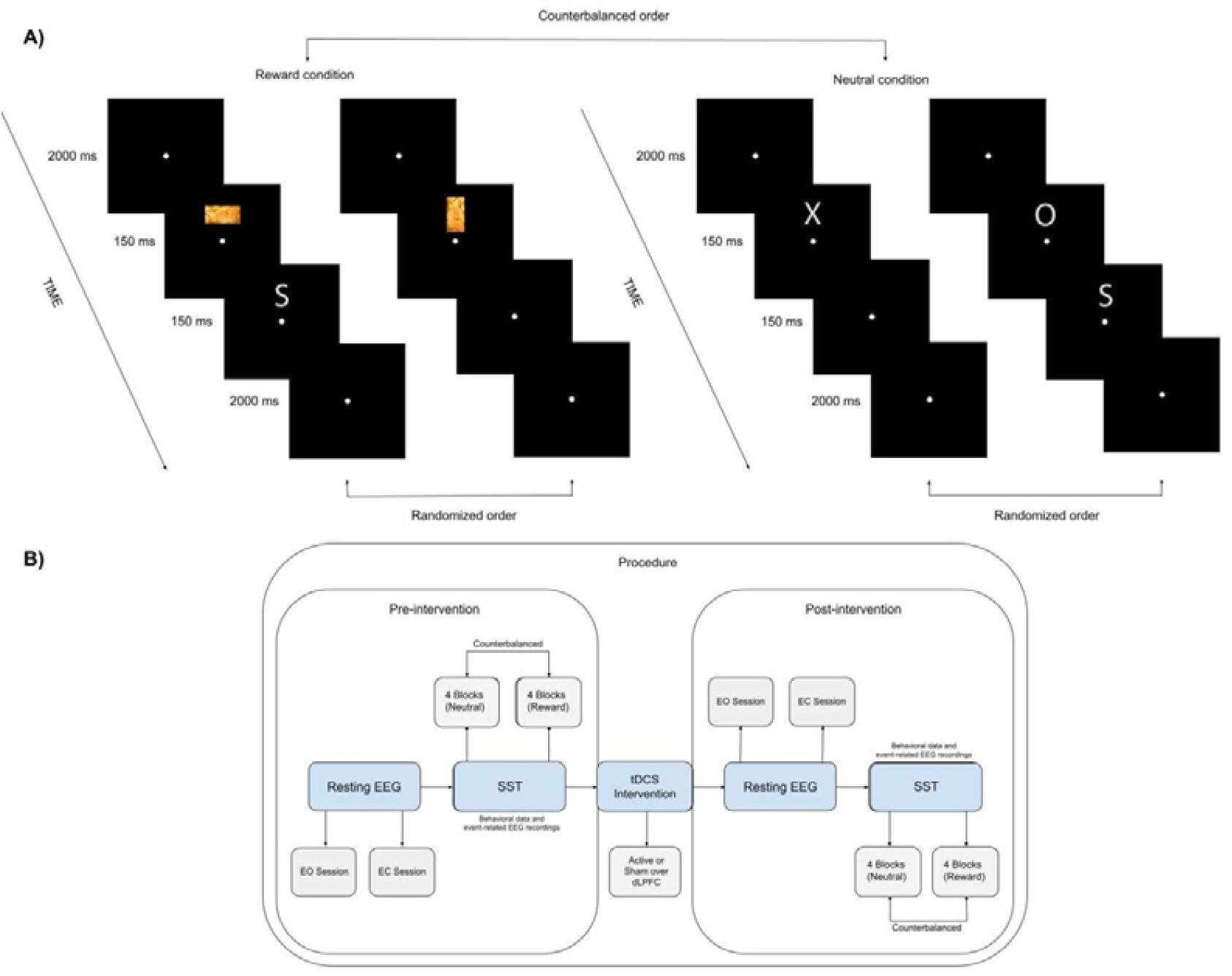
A) Based on the counterbalanced order, participants started the Stop Signal Task under either the neutral or reward condition. The initial two trials (on the left) represent the reward condition, where the target dimension (horizontal/vertical) and the palatable food pictures (such as chips, chocolate, cookies, nuts, etc.) were randomly presented. The final two trials (on the right) represent the neutral condition, where the letters “X” and “O” were randomly presented. These trials were sometimes followed by the letter “S” indicating stop trials, where participants should withdraw their initiated responses to non-stop stimuli displayed beforehand. B) The process begins by gathering pre-intervention resting-state EEG data in order to measure frontal alpha asymmetry, which was conducted in two sessions: one with a 5-minute period of eyes open and another with 5 minutes of eyes closed conditions. Subsequently, participants performed the Stop Signal Task, where both the neutral and reward conditions were presented in a counterbalanced order. Following this, either sham or active transcranial direct current stimulation was administered. More details on the intervention can be found in Figure 2. After the stimulation, the same procedure was repeated for the post-modulation assessment, including the resting-state EEG and the Stop Signal Task.

### 2.3. EEG acquisition

Scalp voltages were captured using a 21-channel cap Ag/AgCl electrodes set according to the 10-20 system ^33^. The vertical electrooculography (VEOG) was recorded above and below the left eye, and the horizontal EOG (HEOG) was recorded bipolarly from the outer canthi of both eyes. Sampling rate was set at 512 Hz. EEG was continually recorded aside from when tDCS was applied because switching from the EEG cap to the tDCS cap was necessary.

### 2.4. Frontal alpha asymmetry (FAA)

FAA scores were derived from EEG data recorded during two separate 5-minute resting-state sessions: one with eyes open (EO) and another with eyes closed (EC) both before and after the intervention. Collecting EEG resting-state recordings under these two circumstances facilitates a clearer grasp of the effects of sensory input, internal cognitive functions, and the inherent activity of the brain. This approach contributes to a thorough comprehension of the functional organization of the brain in various states ^34^. Using BrainVision Analyzer 2 (www.brainproducts.com), FAA was calculated based on previous reports ^35^. First, a high-pass filter of 0.5 Hz, low-pass filter of 40 Hz, and 50 Hz notch filter were applied. Subsequently, the first and last 10 seconds of the EEG data was excluded due to artifacts. The data was then segmented into 2-second epochs. Based on VEOG and HEOG electrodes, the epochs for the EO condition were adjusted for ocular artifacts using independent component analysis (ICA). Epochs that still contained artifacts, determined by a 75 μV maximum amplitude +/- relative to the baseline criterion, were excluded. The Power Spectral Density (PSD) was determined using FFT with a 10% Hanning window for the remaining epochs after whole-segment baseline correction. Next, the epochs were averaged, and the mean activity in the alpha frequency band (8-13 Hz) was calculated. Values for the relevant electrodes were exported. Using SPSS 22 ^36^, alpha power was corrected for skew via natural log transform ^35^. Lastly, frontal asymmetry was calculated by subtracting the log transformed alpha at lateral left electrode sites from right electrode sites (i.e., F4-F3 and F8-F7).

### 2.5. Event-related potentials (ERPs)

First, signals were referenced to linked mastoids. Subsequently, following previous studies ^37,38^, EEG data were filtered (offline) using a high cutoff 30 Hz, low cutoff 0.5 Hz, and notch filter of 50 Hz. Data were segmented into epochs ranging from -100 ms to 2600 ms. Ocular corrections were conducted using Independent Component Analysis. Following that, epochs were segmented and they were baseline-corrected with the baseline set at -100 to 0 ms. We conducted a go-signal locked segmentation, then baseline correction followed by artifact rejection (using minimal/maximal allowed amplitude -75 μV/75 μV and marking 200 ms before and after events as bad). We employed a stop-signal locked segmentation, and baseline correction. Next, we computed separate averages for segments corresponding to failed stops and successful stops. The inhibitory ERPs were computed by subtracting the average stop-signal locked activity for failed stops from successful ones. Following a thorough examination of the grand average waveforms, we have identified specific latency intervals for further analysis. For the Stop N2 component, we extracted data from the time window of 166-286 ms, while for the Stop P3 component, data from the 211-271 ms time window was selected for export.

### 2.6. Transcranial direct current stimulation (tDCS)

The aim of brain modulation was to increase the relative activity of the right frontal cortex, which is linked with inhibitory control. A pair of circular sponges (25 cm2) soaked in saline solution, were used to deliver direct electrical current using STARSTIM-8, Neuroelectrics (www.neuroelectrics.com). According to the 10-20 system, the anode was positioned on the right dLPFC (F4) and the cathode on the left dLPC (F3). Unless it is a sham condition, a steady current of 2 mA was applied for 20 minutes ^9^ (Fig. 2). The safety of this parameter has been shown in healthy subjects ^39^.

**Figure 2.**
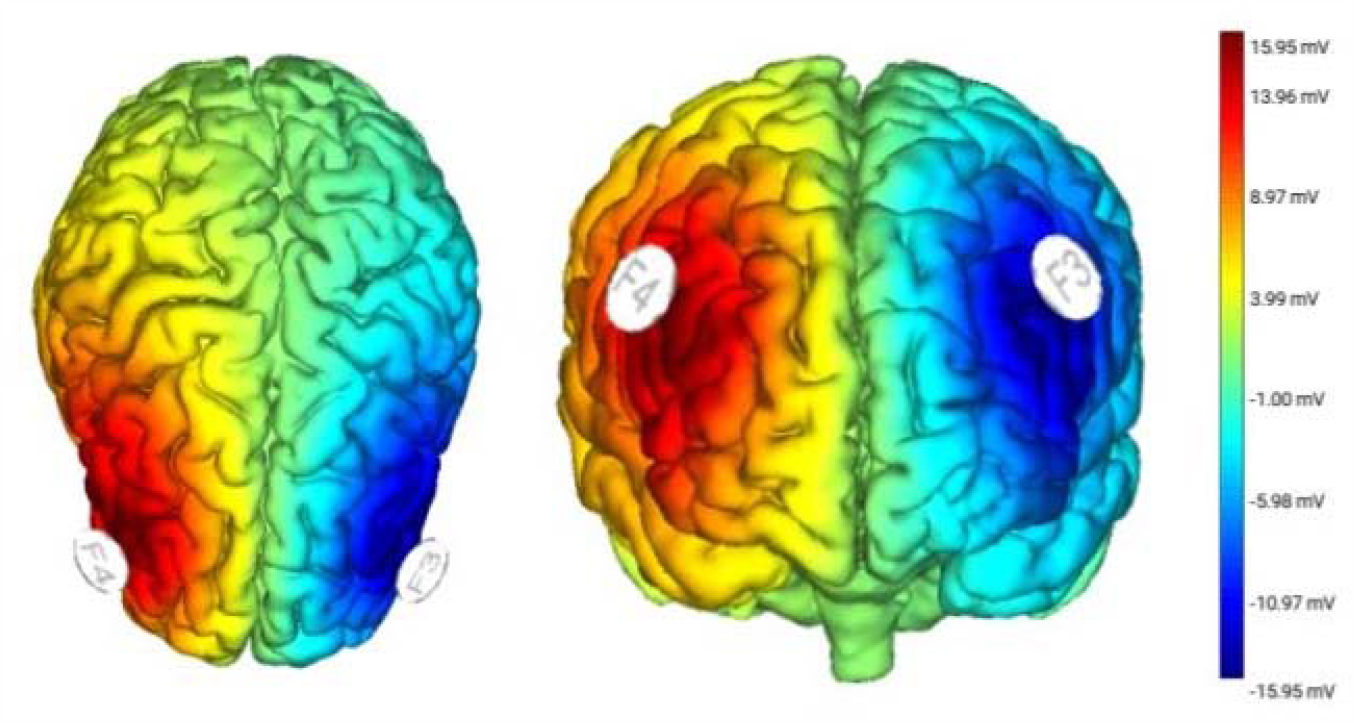
This illustration displays the positioning of the transcranial direct current stimulation electrodes and the measured electric fields viewed from the front and top perspectives. To administer transcranial direct current stimulation, a steady current of 2 mA was passed through two circular sponge electrodes (25 cm2 each), positioned on the scalp over locations F4 (anode) and F3 (cathode) with the use of a saline solution. The maximum strength of the electric field reached 15.95 μV at the anodal electrode.

### 2.7. Procedure

The study was preregistered on Open Science Framework (https://osf.io/6k8up/?view_only=0cfdcc8e92f441e0905a456611a78762). A randomized, triple-blind, sham-controlled design was used in this research, with within-subject (time: pre-/post-assessment, condition: neutral/reward) and between-subject (group: active/sham tDCS) factors. Before and after the neural modulation session, the SST was completed. Participants were seated in a comfortable chair in a dimly lit testing room for the placement of EEG electrodes on the scalp sites after reading the information letter, verifying the inclusion and exclusion criteria, and completing the informed consent form when they arrived at our lab. Subsequently, resting-state EEG data was collected. It was recorded for 10 minutes in total with two blocks of five minutes sessions (EO/EC). They finished the questionnaires after recording the resting-state EEG. They then completed the first phase of the SST prior to the tDCS. EEG was recorded throughout the resting states and computer tasks, but not during the intervention as the cap was changed. Following that, individuals were assigned to either an active or sham tDCS. During the post-modulation evaluation, the same steps—resting-state EEG and the SST— were repeated. The stimulation and pre-/post-assessments were conducted on the same day and lasted approximately five hours.

### 2.8. Statistical analyses

The analyses were conducted using SPSS 22 ^36^ and R ^40^. In addition to employing frequentist statistics, we also conducted a series of Bayesian analyses using JASP ^41^. Upon completing the calculations of the main variables, participants with missing values and outliers beyond 3 standard deviations (SD) from the mean were excluded in case of apparent erroneous data. Subsequently, a repeated measures ANOVA (2x2x2) was employed to examine the hypotheses. The electrophysiological variables investigated included FAA (EO/EC), the Stop N2, the Stop P3, in addition to the behavioral indices of inhibitory control: SSRTs. We also examined the correlation between baseline (pre-modulation) FAA and behavioral and brain activity indices of inhibitory control. For all frequentist statistical analyses, the significance level was set at 0.05 and for the null results, Bayesian approach was used with Bayes factor 01 (BF_01_), which is in favor of null hypotheses (H_0_) over alternative hypotheses (H_1_). More specifically, anecdotal evidence is characterized by a score ranging from 0.33 to 1. A score below 0.33 suggests significant support for H_1_ ^42,43^. The results of Bayesian and exploratory analyses can be found in the supplementary materials.

## 3. Results

### 3.1. Did the tDCS have an effect on resting-state frontal alpha asymmetry?

In order to examine the impact of the tDCS on resting-state FAA scores, a repeated measures ANOVA with time (pre-/post-intervention) as the within-subjects factor and group (active/sham tDCS) as the between-subject factor was conducted. The results of the analysis revealed no significant interaction between the time and group factors concerning FAA. Table 1 shows the details. These results were also supported by Bayesian statistics. More specifically, all reported BF_01_ values in Tables 4, 5, 6, and 7 in the supplementary materials consistently demonstrate substantial evidence in favor of the null hypothesis (BF_01_ > 0.3).

**Table 1.** Results of FAA Models.

| Variables | Df | F | P | $\eta_p^2$ |
| --- | --- | --- | --- | --- |
| FAA F4-F3 (EO) (N = 59) |  |  |  |  |
| Group | 1 | 3.64 | 0.058 | 0.030 |
| Time | 1 | 0.39 | 0.530 | 0.003 |
| Group x Time | 1 | 0.28 | 0.594 | 0.002 |
| Error | 114 |  |  |  |
| FAA F4-F3 (EC) (N = 59) |  |  |  |  |
| Group | 1 | 1.22 | 0.271 | 0.010 |
| Time | 1 | 1.27 | 0.261 | 0.010 |
| Group x Time | 1 | 0.49 | 0.484 | 0.004 |
| Error | 114 |  |  |  |
| FAA F8-F7 (EO) (N = 62) |  |  |  |  |
| Group | 1 | 0.00 | 0.951 | <0.001 |
| Time | 1 | 0.00 | 0.933 | <0.001 |
| Group x Time | 1 | 0.29 | 0.588 | 0.002 |
| Error | 120 |  |  |  |
| FAA F8-F7 (EC) (N = 59) |  |  |  |  |
| Group | 1 | 0.03 | 0.849 | <0.001 |
| Time | 1 | 3.58 | 0.060 | 0.030 |
| Group x Time | 1 | 0.08 | 0.769 | <0.001 |
| Error | 114 |  |  |  |

Note: Participants with missing values and outliers (based on 3 standard deviations from the mean) were excluded from the analyses.

We also conducted a correlation analysis to further explore the relationship between trait FAA and both behavioral and brain activity indicators of inhibitory control. In the reward condition, we observed a significant negative correlation between trait FAA F4-F3 (EO) and two key brain activity indices of inhibitory control, namely the Stop N2 (r = -0.46, p < 0.01) and P3 (r = -0.30, p < 0.05). For further information, refer to Table 8 (in the supplementary materials).

### 3.2. Did the tDCS have an effect on the behavioral indices of inhibitory control?

To assess the effects of the tDCS intervention on inhibitory control, we conducted a repeated measures ANOVA using time (pre-/post-intervention) and condition (neutral/reward) as within-subjects factors, and group (active/sham tDCS) as the between-subject factor on SSRT scores. The results were presented in Figure 3, and inferential statistics are shown in Table 2. We found no significant effect of tDCS on the SSRT. However, there was a significant main effect of time on the SSRT and it was supported by Bayesian statistics as well (F(1, 232) = 6.52, p = 0.011, η_p_^2^ = 0.004, BF_01_ < 0.3). These findings indicate that inhibitory control decreases as time progresses, irrespective of the condition or group. Besides time model, time + group (BF_01_ < 0.3), time + condition (BF_01_ < 0.3), and time + condition + group (BF_01_ < 0.3) models were also in favor of H_1_. For more information, please see Table 9 (in the supplementary materials).

**Table 2.** Results of the SST.

| Variable(s) (N = 58) | Df | F | p | $\eta_p^2$ |
| --- | --- | --- | --- | --- |
| SSRT |  |  |  |  |
| Time | 1 | 6.52 | 0.011* | 0.028 |
| Condition | 1 | 0.99 | 0.319 | 0.004 |
| Group | 1 | 3.82 | 0.051 | 0.016 |
| Time x Condition | 1 | 0.18 | 0.657 | <0.001 |
| Time x Group | 1 | 0.19 | 0.670 | <0.001 |
| Condition x Group | 1 | 1.10 | 0.294 | 0.004 |
| Time x Condition x Group | 1 | 0.02 | 0.883 | <0.001 |
| Error | 232 |  |  |  |

Note: Participants with missing values and outliers (based on 3 standard deviations from the mean) were excluded from the analyses.

**Figure 3.**
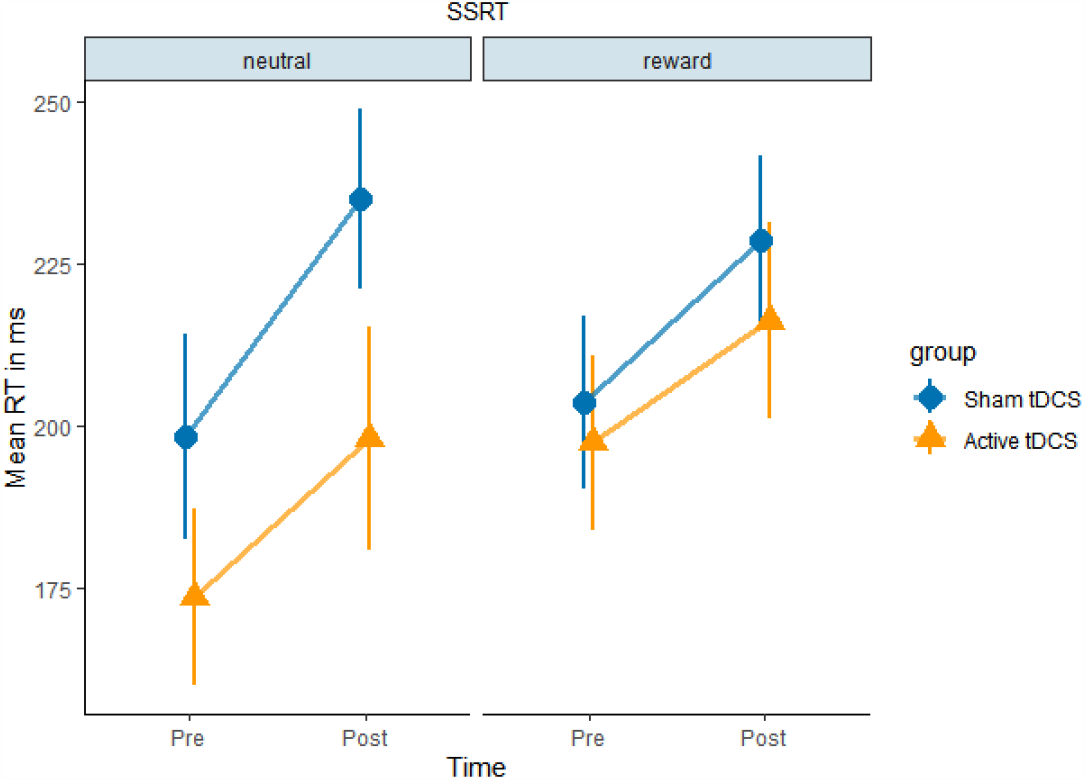
This figure displays the average stop-signal reaction times, considering the factors of time (pre-/post-intervention), condition (neutral/reward), and group (sham/active transcranial direct current stimulation). The error bars indicate standard errors. It illustrates that inhibitory control decreased from pre-assessment to post-assessment, regardless of condition and group factors. It is important to note that longer stop-signal reaction times represent decreased inhibitory control.

### 3.3. Did the tDCS have an effect on the brain activity indices of inhibitory control?

In order to evaluate the impact of the tDCS intervention on the brain activity measures of inhibitory control, we performed a repeated measures ANOVA. This analysis included time (pre-/post-intervention) and condition (neutral/reward) as within-subjects factors, while group (sham/active tDCS) was the between-subject factor. The focus was on the Stop N2 and Stop P3 components. The interaction effect of time, condition, and group on the indices was found to be insignificant. They were displayed in Figure 4 and Figure 5, respectively. All reported models in Bayesian approach were in favor of H_0_ (BF_01_ > 0.3). For further information, please see Tables 10 and 11 in the supplementary materials. However, there is a statistically significant effect of the interaction between time and group on the Stop P3: F(1, 57) = 7.18, p = 0.010, η_p_^2^ = 0.112, as illustrated in Figure 6. More detailed results can be found in Table 3.

**Table 3.** Results of ERP models.

| Variable(s) | Df | F | p | $\eta_p^2$ |
| --- | --- | --- | --- | --- |
| Stop N2 at 166-188 ms. (F4) (N = 59) |  |  |  |  |
| Time | 1 | 1.58 | 0.214 | 0.027 |
| Condition | 1 | 0.27 | 0.602 | 0.005 |
| Group | 1 | 1.46 | 0.231 | 0.025 |
| Time x Condition | 1 | 1.75 | 0.191 | 0.030 |
| Time x Group | 1 | 0.25 | 0.613 | 0.005 |
| Condition x Group | 1 | 0.14 | 0.704 | 0.003 |
| Time x Condition x Group | 1 | 0.42 | 0.517 | 0.007 |
| Error | 57 |  |  |  |
| Stop P3 at 211-271 ms. (Cz) (N = 59) |  |  |  |  |
| Time | 1 | 0.97 | 0.327 | 0.017 |
| Condition | 1 | 1.44 | 0.234 | 0.025 |
| Group | 1 | 3.05 | 0.086 | 0.051 |
| Time x Condition | 1 | 7.18 | 0.010* | 0.112 |
| Time x Group | 1 | 0.12 | 0.726 | 0.002 |
| Condition x Group | 1 | 0.05 | 0.818 | 0.001 |
| Time x Condition x Group | 1 | 0.08 | 0.775 | 0.001 |
| Error | 57 |  |  |  |

Note: Participants with missing values and outliers (based on 3 standard deviations from the mean) were excluded from the analyses.

**Figure 4.**
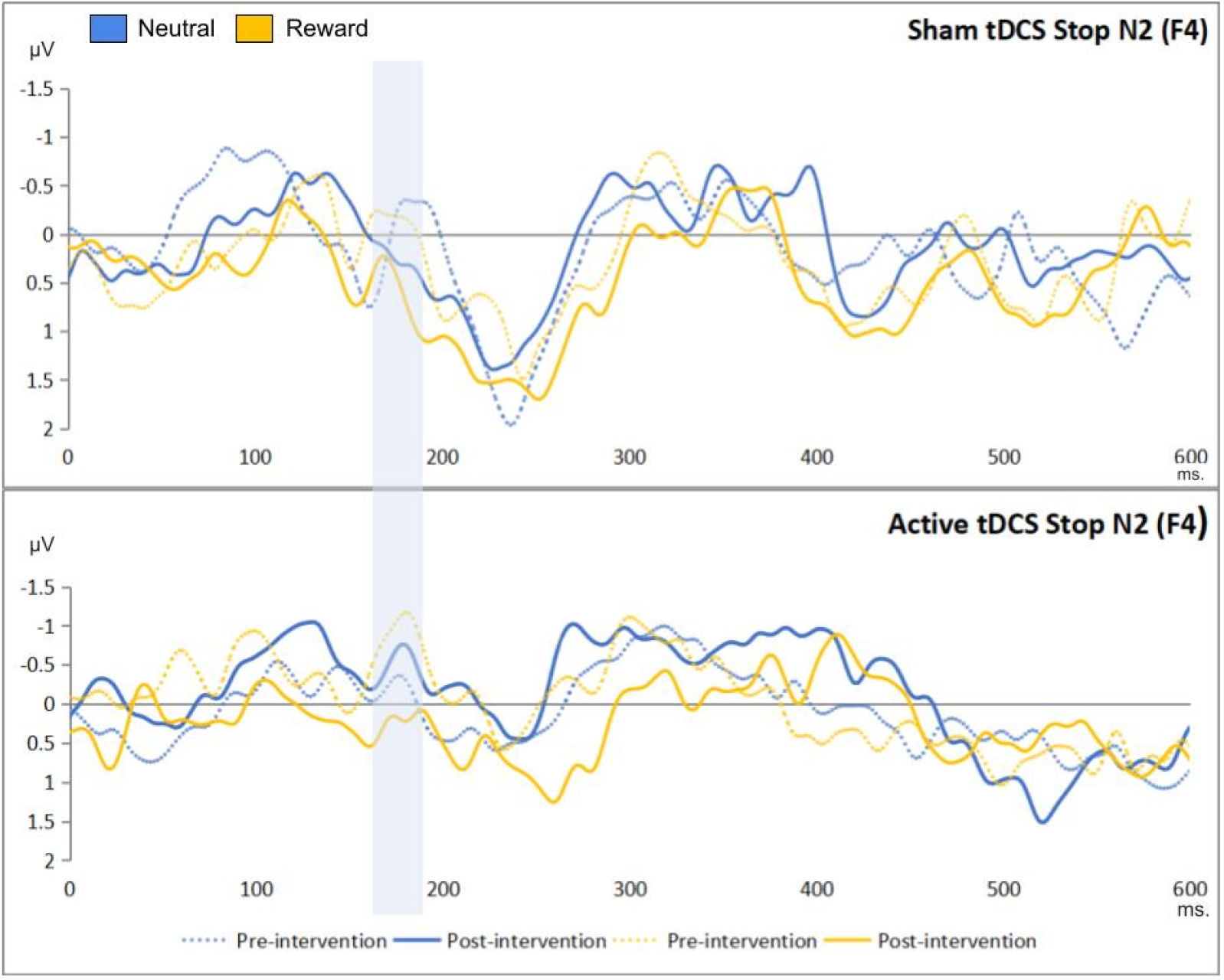
This figure shows the stop-signal locked event-related potentials during the stop signal task, effects of time, condition, and group on the Stop N2 (166-286 ms), based on successful inhibitions minus failed ones. The x-axes represent the time in milliseconds; the y-axes represent the Stop N2 scores in microvolts.

**Figure 5.**
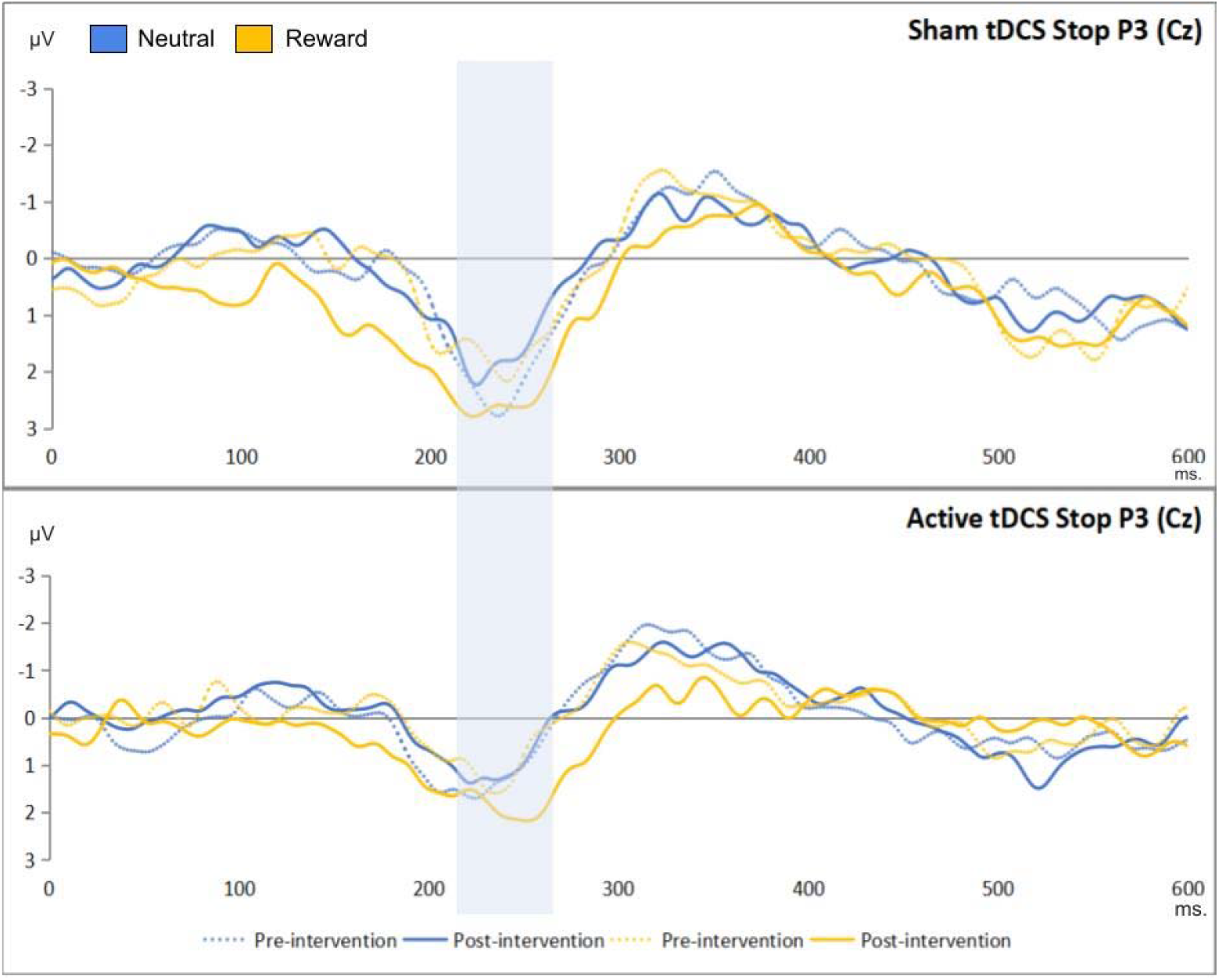
This figure shows the stop-signal locked event-related potentials during the stop signal task, the effects of time, condition, and group on the Stop P3 (211-271 ms), based on successful inhibitions minus failed ones. The x-axes represent the time in milliseconds; the y-axes represents the Stop P3 scores in microvolts.

**Figure 6.**
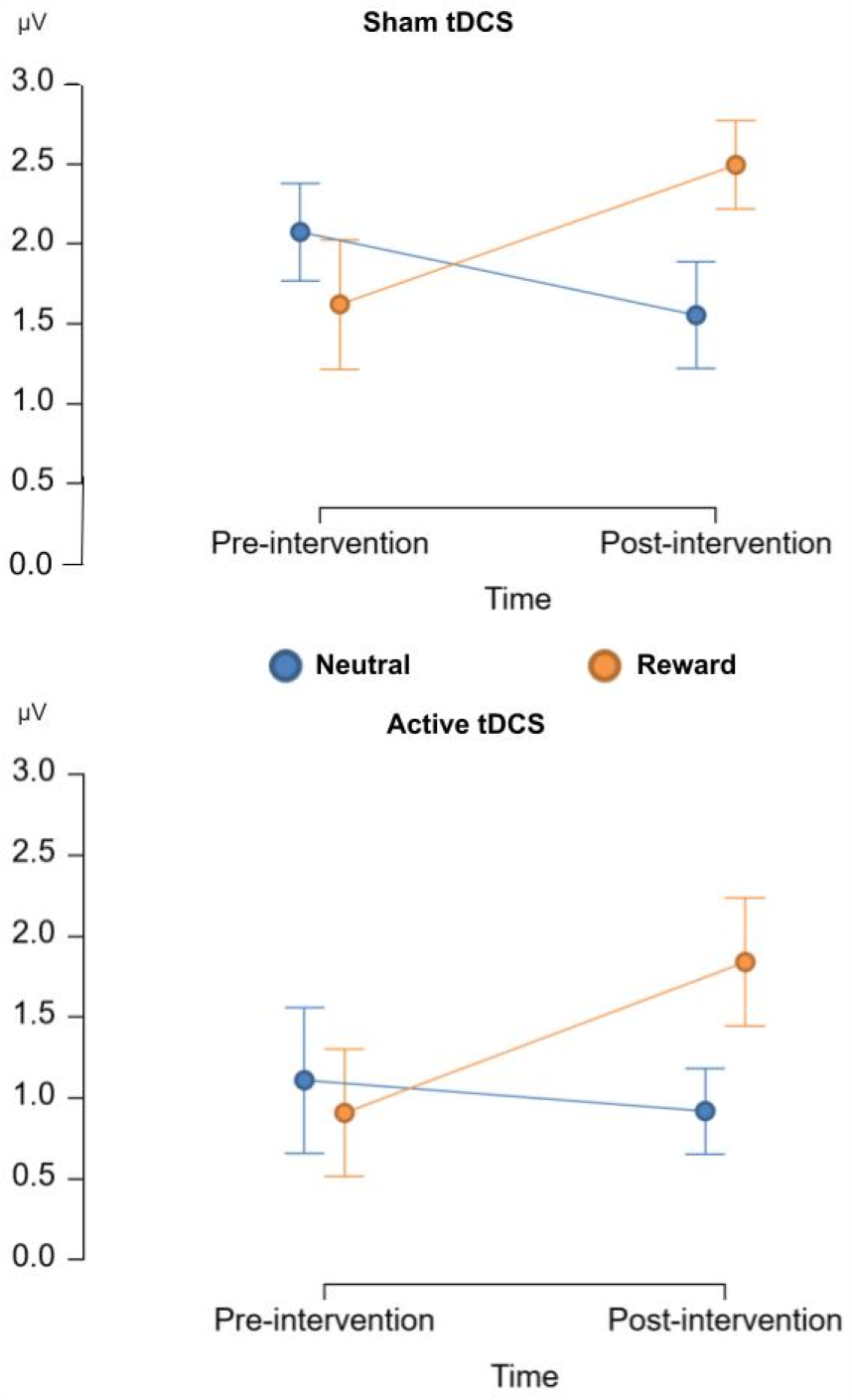
The figure shows the exact effect of time and group interaction on the Stop P3. The x-axes represent the time factor; the y-axes represent the Stop P3 in microvolts. The error bars illustrates standard errors. It indicates a higher inhibitory activity in the reward condition compared to the neutral condition.

## 4. Discussion

Our study aimed to explore the potential effect of tDCS on resting-state FAA, a promising transdiagnostic neuromarker for inhibition-related clinical conditions, and to investigate whether the effect of tDCS on inhibitory control is mediated by its influence on FAA. Contrary to our initial hypothesis, we did not observe any significant connections between tDCS and FAA, and there were no noticeable impacts on the behavioral and brain activity indicators of inhibitory control. Interestingly, greater right frontal brain activity compared to the left (indicating lower frontal alpha asymmetry) was found to be correlated with reduced early-onset inhibition (as evidenced by Stop N2), yet it was associated with heightened late onset inhibitory activity (as indicated by Stop P3). Furthermore, a noteworthy trend emerged wherein participants’ SSRT showed a decline across all conditions and groups, with the concurrent rise in inhibitory brain activity, particularly prominent in the reward condition, as indicated by the stop-signal locked P3 component.

Previous research has suggested a potential influence of tDCS on the asymmetry of frontal brain activity ^22^. However, these measures could be considered indirect due to the absence of comprehensive research into the direct impact of tDCS on the primary target, frontal asymmetry. One candidate measure for frontal asymmetry is the FAA. We conducted a pilot study and found that the smallest effect of tDCS on FAA, with direct measurement of resting states, could be reached with 30 participants. Nevertheless, our study did not find any support for the effect of tDCS on FAA. The intricate nature of tDCS effects on brain activity, combined with the inherent individual variability in brain anatomy, may have contributed to the absence of significant findings in our research. More specifically, the path of electric currents during tDCS involves a multifaceted journey through different tissues and layers, with varying degrees of resistance and susceptibility to neural alteration. Each participant’s unique brain structure, including factors like skull thickness and sulcal depth, could have resulted in divergent responses to tDCS ^44,45 46^. Moreover, factors related to electrode positioning and properties may have played a role in shaping the outcomes. The size, shape, and conductivities of the electrodes, as well as the use of gels and saline solutions, may have influenced the distribution and intensity of the electric field ^47^. Furthermore, frontal tDCS is known to produce more variable electric fields compared to other types of tDCS ^48^, adding further complexity to the neural adjustment processes. While our findings challenge the direct impact of tDCS on FAA, they underscore the importance of considering individual differences and optimizing stimulation protocols in future research.

Similarly, our investigation into the effect of tDCS on the various indices of inhibitory control yielded no significant results. The impact of tDCS on cognition and behavior is notably variable, challenging the notion of a polarity-specific influence. While tDCS is theoretically expected to increase excitability under the anode and decrease it under the cathode, the actual cognitive and behavioral effects are far more complicated. Interestingly, some studies have even reported facilitatory effects associated with stimulation under the cathode, possibly attributed to noise reduction in specific networks, leading to improved performance ^49^. Alternatively, cathodal tDCS might inhibit a particular function as well, leading to enhanced performance in specific tasks, like faster reaction times ^50^. Numerous factors contribute to these contradictory outcomes. For example, the effects of prefrontal tDCS heavily depend on the state of the targeted neural network ^51^. In the online paradigm, tDCS influences networks already engaged in the task, while in the offline paradigm, it modifies neural activity beyond the stimulation period. Understanding these state-dependent effects is crucial for cognitive and behavioral studies, as factors like fatigue, task knowledge, and network connectivity can significantly influence the baseline neural state. Moreover, a recent meta-analysis ^52^ extensively examined the effects of tDCS on go/no-go and SST performance measures of inhibitory control in single-session designs. The findings revealed that the placement of electrodes significantly influences the outcomes. Specifically, tDCS targeting the right inferior frontal gyrus demonstrated a medium effect size, whereas a stimulation site over the dLPFC region showed an overall null effect. Variation in results was attributed to the positioning of the return electrode, with extracephalic placement differing from various positions across the head. These results underscore the critical importance of precisely targeting specific brain regions in tDCS research to achieve desired and consistent outcomes. Another plausible explanation could be attributed to the sample characteristics. Research on binge eating ^21–23^ suggests that tDCS primarily affects inhibitory control in samples with considerable room for improvement. This hypothesis proposes that individuals with relatively weaker inhibitory control at baseline may experience more pronounced enhancements following tDCS intervention. Conversely, in healthy individuals, the influence of tDCS may not predominantly target inhibitory activity but rather attentional bias ^53^.

Our correlation analysis revealed a significant connection between inhibitory brain activity during reward conditions and trait frontal brain asymmetry, as indexed by resting-state FAA before the neurostimulation. Precisely, greater right-sided frontal brain activity compared to the left side was linked to reduced initial inhibitory activity (Stop N2), likely emanating from the inferior frontal gyrus. However, it was also associated with heightened subsequent inhibitory control (Stop P3), which is thought to originate from the superior frontal gyrus. The brain is organized into functional networks, and the interaction between different regions is crucial for complex processes. The observed correlation might suggest a dynamic interplay between the inferior and superior frontal gyri in managing inhibitory control. Higher activity in one region may trigger or facilitate inhibitory control processes in another region. Furthermore, the higher right frontal brain activity might signify a compensatory mechanism. When early-onset inhibitory activity originating from the inferior frontal gyrus is compromised, the brain might engage the superior frontal gyrus to enhance inhibitory control at a later stage. Alternatively, the nature and demands of the task being performed could influence how inhibitory control is exerted. Based on that, the brain’s inhibitory processes might operate differently at different stages of a task such as at early-onset inhibitory activity, associated with the inferior frontal gyrus, and late onset inhibitory control involving the superior frontal gyrus to achieve optimal inhibitory control. It is important to note that the interpretation provided is speculative and would need to be validated through empirical research and neuroimaging studies. The brain is very complex, and multiple factors could contribute to the observed dissociation between brain activity and inhibitory control.

Lastly, based on the SST results, a notable reduction in inhibitory control was observed as time progressed, which aligns with expectations due to factors like tiredness and fatigue. However, the ERP results revealed an intriguing pattern in the reward context, where enhanced inhibitory activity in the brain (indexed by the stop P3) was observed as time progressed, relative to the neutral context. These results suggest that despite the absence of a significant time and condition interaction concerning SSRTs, the post-test reward block posed a stronger inhibitory challenge. As a consequence, participants displayed an adapted response marked by increased inhibition-related activity in the brain. This adaptive neurophysiological response may reflect the brain’s capacity to dynamically adjust and allocate cognitive resources in response to varying levels of inhibitory demand. Consistent with our findings, prior studies have also observed increased N2 and P3 amplitudes during food-specific trials ^54^. In contrast, another study demonstrated that P3, but not N2, significantly decreased in obese participants across all Go/No-go task conditions compared to normal-weight controls, suggesting that P3 might serve as a more critical biomarker of inhibitory control deficits ^55^. However, it is important to note that variations in stimuli, paradigms, component timescales, and ERP analyses present challenges in synthesizing results across the existing literature. The diversity in methodologies utilized calls for caution in drawing definitive conclusions from the available evidence. Future research could benefit from standardized protocols and methodologies to address these complexities, enabling more robust comparisons and a deeper understanding of the neurophysiological underpinnings of inhibitory control in various populations.

Despite the valuable insights obtained from this research, several limitations require careful consideration. It is important to acknowledge that individual variations in brain anatomy might have contributed to the lack of significant effects observed. Additionally, the effectiveness of tDCS depends on various factors, including stimulation parameters (such as duration, intensity, and electrode placement) and target locations, as discussed. Although the chosen intervention parameters were based on existing literature and logical reasoning, alternative stimulation settings might have yielded different outcomes. Using neuroimaging techniques, such as functional magnetic resonance imaging (fMRI), could provide valuable insights into the neural changes induced by tDCS. To gain a deeper comprehension of the impact of tDCS on FAA in relation to inhibitory control, it would be beneficial to carry out studies involving more diverse participant samples. Our Bayesian analyses indicated that there is support for H1 for behavioral index of inhibitory control: SSRT. Therefore, the sample and effect sizes can be considered as a limitation. More specifically, using relatively a small-sample (N = 10) pilot study to identify the effect size of interest can yield extreme results. This can lead to an overestimation of the effect size, which, in turn, can result in an under-powered study. Furthermore, the effect size was calculated for the main effect of tDCS on the primary outcome, FAA, which is different than more complex models in the study. By carefully considering individual brain differences, electrode-related factors, and tDCS parameters, and sample characteristics researchers can enhance the reproducibility and reliability of tDCS studies on prefrontal cortex and inhibitory control.

## Conclusion

In our study, we aimed to explore the effects of transcranial direct current stimulation on frontal brain asymmetry and whether the effects of the transcranial direct current stimulation intervention on inhibitory control-related indices could be attributed to changes in frontal alpha asymmetry. The results indicated that transcranial direct current stimulation did not yield significant effects on frontal alpha asymmetry. However, we made a notable discovery: there was a dissociation between baseline frontal alpha asymmetry and brain activity indices of inhibitory control. Furthermore, the stop signal reaction times consistently increased over time, suggesting reduced inhibitory control. Intriguingly, we also observed a simultaneous increase in inhibitory control-related brain activity, particularly in the post-stimulation reward condition. This finding probably indicates a dynamically adaptive response in the presence of food stimuli. Consequently, although the tDCS yielded negative outcomes, our study reveals that frontal alpha asymmetry still holds the potential to predict neural responses associated with within-task events. This intricate interplay underscores the dynamic relationship between frontal alpha asymmetry and inhibitory processes.

## Notes

### Competing Interest Statement

The authors have declared no competing interest.

### Summary of Updates

Bayesian statistics was added to the manuscript.

